# Loss of FAM111B protease mutated in hereditary fibrosing poikiloderma syndrome negatively regulates telomere length

**DOI:** 10.1101/2023.01.22.525054

**Authors:** Maciej Kliszczak, Daniela Moralli, Julia D. Jankowska, Paulina Bryjka, Lamia Subha Meem, Tomas Goncalves, Svenja S. Hester, Roman Fisher, David Clynes, Catherine M. Green

**Author notes:** Oxford Drug Discovery Institute, University of Oxford, Old Road Campus, Oxford OX3 7FZ, United Kingdom (Maciej Kliszczak). Vienna BioCenter, University of Vienna, 1030 Vienna, Austria (Julia D. Jankowska). The Jenner Institute, University of Oxford, Old Road Campus, Oxford OX3 7DQ, United Kingdom (Paulina Bryjka). These authors contributed equally to this work (Julia D. Jankowska, Paulina Bryjka, Lamia Subha Meem).

## Abstract

Hereditary fibrosing poikiloderma (HFP) is a rare human dominant negative disorder caused by mutations in the *FAM111B* gene that encodes a nuclear trypsin-like serine protease. HFP patients present with symptoms including skin abnormalities, tendon contractures, myopathy and lung fibrosis. We characterised the cellular roles of human FAM111B using U2OS and MCF7 cell lines and report here that the protease interacts with components of the nuclear pore complex. Loss of *FAM111B* expression resulted in abnormal nuclear shape and reduced telomeric DNA content suggesting that FAM111B protease is required for normal telomere length; we show that this function is independent of telomerase or recombination driven telomere extension. Even though *FAM111B*-deficient cells were proficient in DNA repair, they showed hallmarks of genomic instability such as increased levels of micronuclei and ultra-fine DNA bridges. Interestingly, FAM111B variants, including mutations that cause HFP, showed more frequent localisation to the nuclear lamina suggesting that accumulation of mutant FAM111B at the nuclear periphery may drive the disease pathology.

## Introduction

Hereditary fibrosing poikiloderma (HFP also known as POIKTMP, OMIM: 615704) is a rare dominant negative syndrome that affects multiple tissues throughout the body (reviewed in [1-5]). Initial disease symptoms appear very early in childhood and include skin abnormalities such as pokiloderma, photo-induced lesions, loss of facial hair and baldness. Many of these symptoms persist until adulthood. These phenotypes are accompanied by reduced sweating, heat intolerance and swelling of the extremities. Tendon contractures require medical attention to improve mobility of young HFP patients. As the affected individuals progress into adulthood other symptoms become more apparent. These include muscular wasting, digestive dysfunction (exocrine pancreatic insufficiency), respiratory obstruction (lung fibrosis) and pancreatic cancer. Furthermore, fibrotic lesions in the respiratory tract and skin can be detected in the in the affected individuals [5-19].

HFP is caused by mutation of a single allele of the *FAM111B* gene that is located on chromosome 11 (11q12.1). *FAM111B* encodes a 734aa protein with a trypsin-like protease domain at its C-terminus. It is 42% identical to the homologous serine protease FAM111A which is mutated in Kenney-Caffey and Gracile Bone Dysplasia syndromes, characterised by severe skeletal abnormalities [20]. FAM111A localises to DNA replication forks via a PIP-box mediated interaction with PCNA, where it processes DNA-protein crosslinks in response to replication stress [21]. Like FAM111A, the FAM111B protease has a conserved catalytic triad, (residues H490, D544, S650) and shows protease activity *in vitro* [22]. Even though FAM111B does not contain a PIP box sequence, similarly to FAM111A it interacts with PCNA and the RFC complex [22]. FAM111B also possesses a centrally located FAM111B-specfic domain (112aa) of unknown function. The substrates and function of FAM111B, and thus the mechanism by which its mutation causes disease are currently unknown.

FAM111B is a nuclear protein expressed in tissue specific manner (brain, lungs, gastrointestinal tract, liver, pancreas, reproductive tissue, skin, bone marrow and lymphoid tissue) with peak cellular concentration during S phase of the cell cycle [23-25]. Arrest of the cell cycle outside S phase leads to down-regulation of FAM111B mRNA and protein levels [26, 27]. Up until now, eight different FAM111B disease causing mutations have been described (reviewed in [1-5]). These affect single amino acids and cluster around two different regions of FAM111B, either in the protease domain (Y621D, T625N, R627G, S628N and S628G) or in the region of unknown function, very close to the FAM111B-specific domain (F416S, ΔK421 and Q430P). Given the autosomal dominant inheritance pattern of the disease, it is likely that these mutations result in the acquisition (or potentially loss) of protein function that has particularly harmful consequences for instance in cells of the connective tissue.

Telomeres are DNA sequences found at the end of chromosomes, which in humans consist of multiple TTAGGG repeats [28]. The shortening of telomeres is a naturally occurring process that coincides with human aging, and re-expression of telomerase in human cells result in unlimited proliferation capacity [29]. Short telomeres are recognised as DNA double strand breaks (reviewed in [30]) and activate p53 pathway leading to apoptosis or senescence (reviewed in [31]). Such an arrangement allows for a strict control of somatic cell’s proliferation capacity and suppresses duplication of cells with abnormally short telomeres (reviewed in [32]). Premature telomere shortening is detrimental for normal cellular function and has been linked to multiple human disease including lung fibrosis (reviewed in [33] and [34-38]).

Here we show that cells lacking FAM111B protease show hallmarks of genomic instability and have short and abnormal telomeres. Telomere lesions were induced in both telomerase-positive and negative cells strongly suggesting that FAM111B was not directly involved in either telomerase-or recombination-dependent telomere extension mechanisms. Interestingly, FAM111B protease could be detected at the nuclear periphery in the proximity of the nuclear envelope and interacted with components of the nuclear pore complexes. Mutant FAM111B, including HFP-linked amino acid substitutions, localised more frequently to the nuclear lamina suggesting that accumulation of FAM111B at the nuclear periphery might drive disease pathology.

## Results

### FAM111B protease localises to the nuclear lamina and interacts with nuclear pore complexes

Staining of human cells with FAM111B antibodies revealed that the protease can be found in both cytoplasm and nucleus; however, the majority of FAM111B resided in the latter compartment (but not in nucleoli) (Figure 1A). The modulation of FAM111B protein levels, dependent on cell cycle position was very apparent and consistent with previously published reports [22, 24, 25] (Figure 1A and B). Fractionation of U2OS cells showed that FAM111B was soluble with a fraction of the protease also bound to insoluble components of the cell such as chromatin or cellular membranes (Figure 1B). To determine whether the immobilised fraction of the FAM111B accumulated in a particular nuclear compartment, we analysed FAM111B staining in U2OS cells using high resolution microscopy. Prior to analysis, soluble and proteins weakly bound to the structural components of the cell (such as cytoskeleton or chromatin etc.) were removed by detergent extraction and the remaining cellular material stained with antibodies against FAM111B and Lamin A/C. FAM111B staining in these conditions revealed a clear three-dimensional organization inside the nucleus (Figure 1C and Movie 1A) again no nucleolar staining was observed. Interestingly, FAM111B also localised to the nuclear periphery where it either interacted with or was found embedded in the nuclear lamina (Figure 1C, white arrow heads, Movie 1B). This suggested that FAM111B interacted with the lamin network. To test whether FAM111B directly interacted with lamin proteins, we characterised FAM111B binding partners using mass spectrometry. Analysis of the FAM111B interactome revealed that FAM111B did not form complexes with lamins but instead with components of the nuclear pore complexes (NPCs) (Figure 1D and Supplementary 1A). This was consistent with the observed localisation of FAM111B and suggested that FAM111B may be recruited to nuclear periphery through the interaction with components of the NPCs.

**Figure 1.**
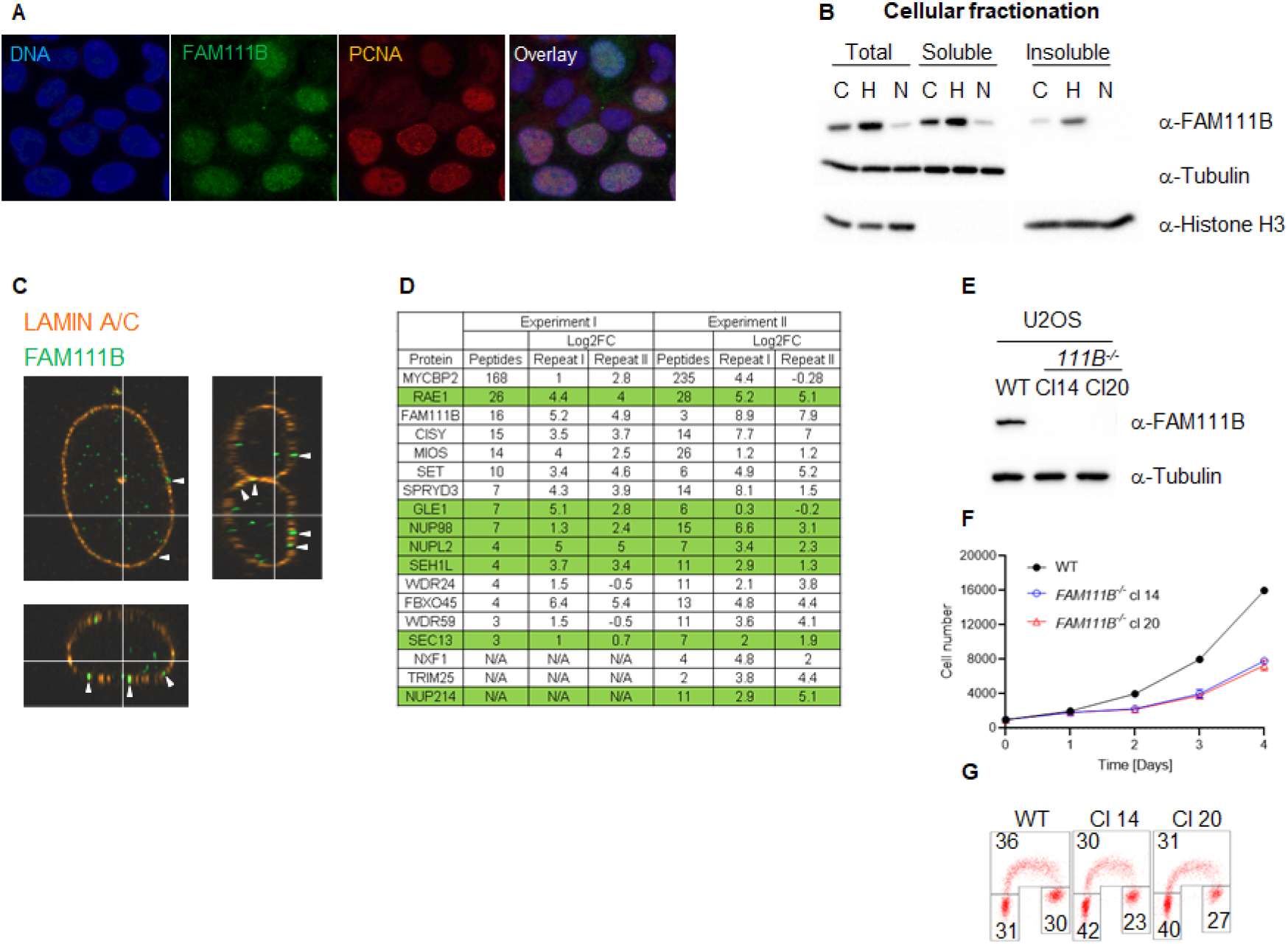
FAM111B localises to the nuclear lamina where it interacts with nuclear pore complexes. A) Images showing FAM111B and PCNA staining in U2OS cells. B) Immuno-blots of different cellular fractions of U2OS cells before and after treatment with cell cycle inhibitors hydroxyurea (H, S -phase arrest) and Nutlin-3a (G1 and G2/M arrest). C) Images of detergent extracted U2OS cells stained with FAM111B and Lamin A/C antibodies. D) FAM111B interacting partners detected by mass spectrometry. E) Immuno-blots showing FAM111B-deficient clones in U2OS cells. F) Growth curves of U2OS wild-type and two FAM111B knockout clones. G) Flow cytometry analysis of the cell cycle distribution in U2OS wild-type and two FAM111B knockout clones.

### Different variants of FAM111B, including HFP mutations cause accumulation of the protease at the nuclear periphery

Localisation of the endogenous FAM111B protease at the nuclear periphery and its interactions with the nuclear pore complexes suggested that this localisation may be essential for the normal function of FAM111B. We wondered whether different variants of FAM111B including mutations found in the HFP syndrome could affect this localisation. We over-expressed different FAM111B variants in U2OS cells, extracted the soluble proteins and stained the cells with FAM111B antibodies (Figure 2A-D). FAM111B wild-type (WT) as well as the <u>p</u>rotease <u>d</u>ead (PD, H490A) variant of the protein showed similar staining pattern to the endogenous FAM111B (Figure 2A and data not shown). Interestingly, in the cells over-expressing HFP variant (Q430P, M1) we could observe a small fraction of the cells where FLAG-FAM111B very strongly accumulated at the nuclear periphery (Figure 2B). Because the HFP mutant variants of FAM111B undergo auto-cleavage into proteolytic fragments (Figure 2E and Supplementary Figure 1A) [22], we investigated whether the resulting fragments had similar potential to localise to the nuclear periphery. The auto-cleavage site in the FAM111A (F334) protease is in FAM111B (F438, Supplementary Figure 3E) [21], and the theoretical molecular weight of the N-terminal fragment after cleavage at this site corresponded to the actual mass detected by immunoblotting (data not shown). We cloned both predicted fragments: N-terminal cleavage fragment (NCF) and C-terminal cleavage fragment (CCF) and over-expressed them in U2OS cells as FLAG-fusion proteins. NCF fragment showed similar localisation to HFP mutation, with strong accumulation at the periphery (Figure 2C) whereas the C-terminal fragment did not express to detectable levels. Cellular fractionation of the cells over-expressing FAM111B with disease relevant HFP mutation showed that the N-terminal auto-cleavage fragment accumulated exclusively in the insoluble fraction of the cell (Figure 2E). Furthermore, we could also observe that FAM111B lacking a centrally located domain (Δ279-386aa) which is not conserved between FAM111A and FAM111B (FAM111B <u>e</u>xtra <u>d</u>omain, ED) caused exclusive accumulation of the FAM111B protease at the nuclear periphery (Figure 2D). Our analysis suggested that FAM111B activity might be transiently required at the nuclear periphery and that accumulation of HFP associated FAM111B variants or truncated forms at this localisation could be potentially pathological.

**Figure 2.**
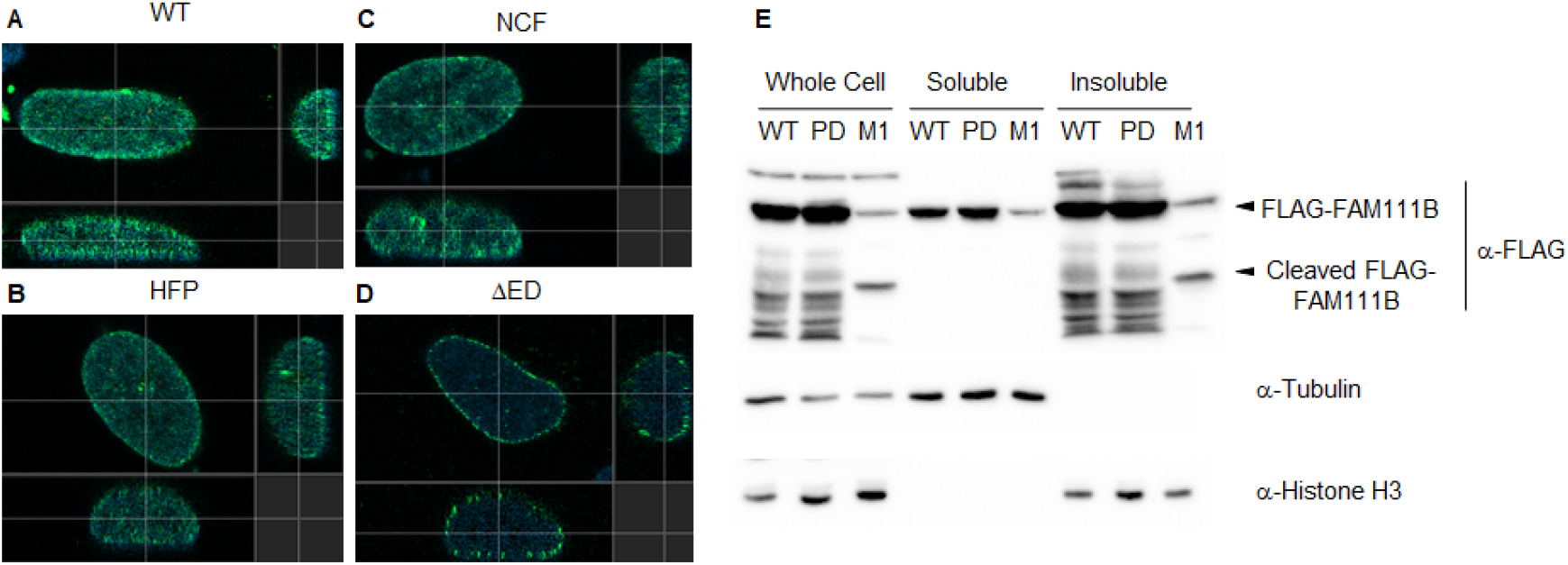
Localisation of mutant FAM111B to the nuclear periphery. A) to D) Images showing localisation of different FAM111B variants: wild-type (WT), Q430P (HFP), N-terminal cleavage fragment (NCF) and ΔED (Δ279-386). E) Immuno-blots of cellular fractions of U2OS cells over-expressing FLAG-tagged FAM111B wild-type (WT), protease dead (PD) and HFP mutant (Q430P, M1).

### FAM111B-deficient cells have abnormal nuclear morphology

Because FAM111B localised to the nuclear lamina we predicted that loss of FAM111B may have a negative effect on nuclear morphology. To test this hypothesis, we employed a reverse genetic approach and manipulated the *FAM111B* gene in human cells (U2OS and MCF7) using Cas9-mediated editing and two independent guide RNAs (Figure 1E and Supplementary Figure 1B). We screened targeted clones through fluorescent in <u>s</u>itu <u>h</u>ybridisation (FISH) and selected those which had a karyotype identical to wild-type cells and no chromosomal abnormalities at the targeted *FAM111B* loci (Supplementary Figure 1E, and data not shown). As previously reported [22, 39-41], loss of FAM111B expression was compatible with cellular viability; however, U2OS cells lacking FAM111B showed slight growth defects characterised by accumulation of cells in G1 phase of the cell cycle (Figure 1F, G and Supplementary Figure 1B, C and D). Staining of *FAM111B*^*-/-*^ cells with antibodies against lamins showed that the *FAM111B*-deficient cells had increased levels of nuclear abnormalities such as abnormal shape (including nuclear blebbing) and doughnut nuclei (Figure 3A and B, Supplementary Figure 1F and G), suggesting that FAM111B function at the nuclear lamina may be required for the maintenance of normal nuclear shape. Similar phenotypes are observed in cells with lamin mutations or other genetic backgrounds caused by deficiencies in the components of the lamina network (reviewed in [42]). We tested whether loss of FAM111B protease expression had any effect on the stability of lamin proteins. We found normal levels of Lamin B1 and only a small change in the Lamin A to C ratio (Figure 3C) in the cells lacking FAM111B, suggesting that the protease is very unlikely to be involved in direct processing or regulation of lamin stability.

**Figure 3.**
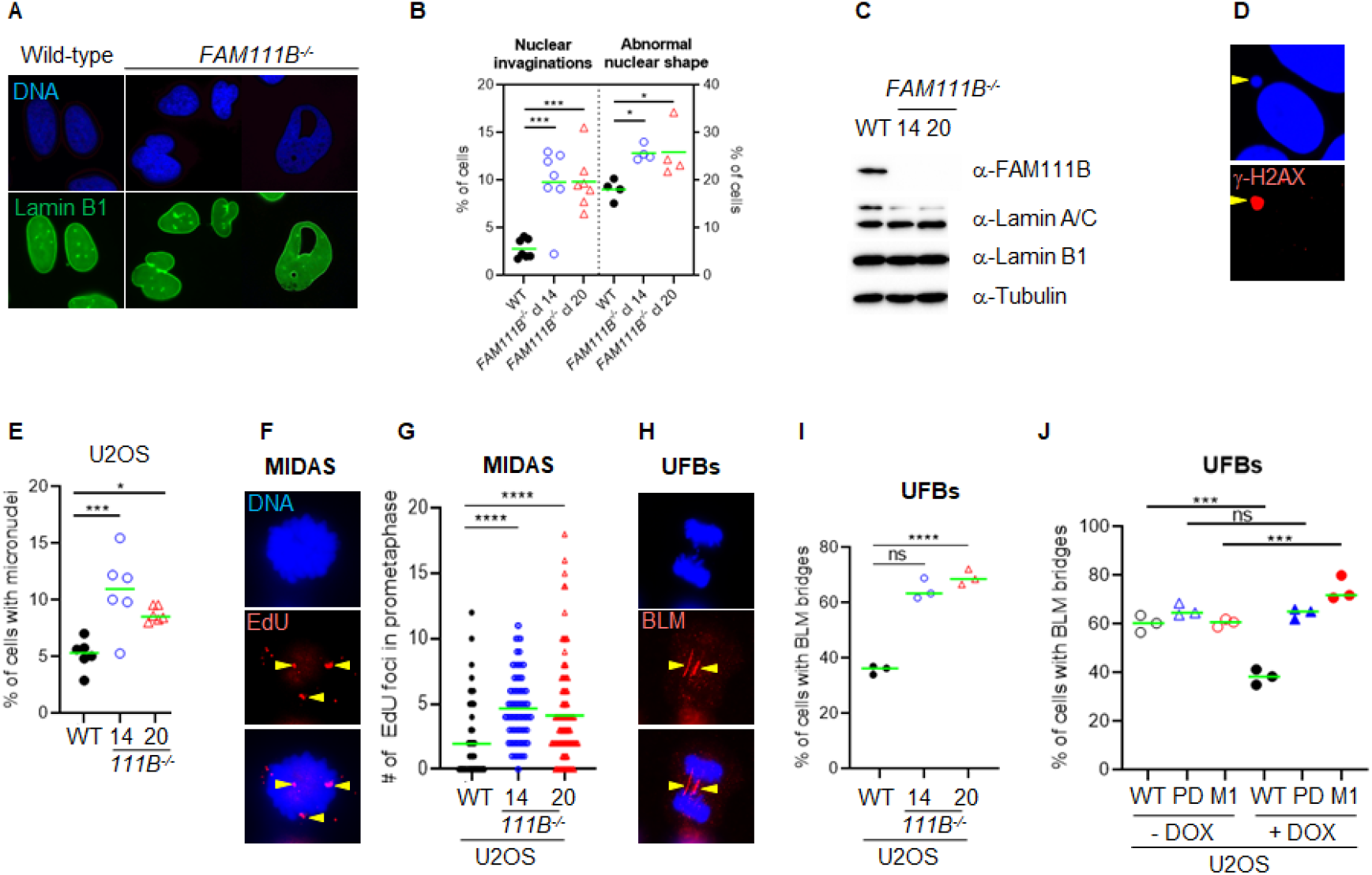
Loss of FAM111B induces abnormal nuclear morphology and spontaneous DNA damage. A) Images showing Lamin B1 staining in U2OS wild-type (WT) and FAM111B-deficinet cells. B) Quantification of nuclear abnormalities in FAM111B knockout cells. C) Immuno-blots showing normal levels of lamin proteins in the cells lacking FAM111B. D) Images of micronuclei. E) Quantification of micronuclei levels in FAM111B-deficient cells. F) Images of MIDAS. G) Quantification of MIDAS in FAM111B knockout cells. H) Images of ultra-fine DNA bridges. I) and J) Quantification of ultra-fine DNA bridges in FAM111B knockout and FAM111B cell lines rescued with either FAM111B wild-type, protease dead (PD) or HFP mutant (Q430P, M1).

### FAM111B is dispensable for DNA repair but loss of FAM111B in U2OS cells increased levels of endogenous DNA damage

The association of FAM111B with nuclear pore complexes suggested a potential role of the protease in the nuclear transport. We sequenced mRNAs isolated from nuclear and cytoplasmic fractions of wild-type and *FAM111B* KO cells and found no major differences in the cellular mRNA content in the absence of FAM111B (data not shown). Consistently, cellular fractionation experiments and immunofluorescence staining showed no measurable differences in the levels of multiple proteins used across this study (data not shown). This suggested that global mRNA and protein transport is not affected by loss of FAM111B expression; however, we cannot rule out the possibility that transport of specific mRNAs or proteins may be affected in the absence of FAM111B. Beside NPCs canonical function as pores, they also serve as hubs for recruitment of chromatin and nuclear factors required for RNA synthesis, DNA repair and telomere maintenance (reviewed in [43]). Nuclear pore complexes participate in maintenance of genome stability through the recruitment of DNA repair intermediates such as double strand breaks and collapsed replication forks [44, 45]. Therefore, we wondered whether similarly to nuclear pore complex mutants [46-48], cells lacking FAM111B could be DNA repair deficient. We treated wild-type and mutant cells with compounds that induce DNA damage and with compounds that interfere with cell cycle, DNA replication or transcription. We analysed survival of wild-type and mutant cells using a resazurin viability assay. Because U2OS cells showed slight growth defect that prevented us to measure cell survival in these experiments, we focused only on MCF7 *FAM111B*-null cells, which were no more sensitive than their wild-type counterparts to multiple drugs tested (Supplementary Figure 2A-J). This indicated that FAM111B was very unlikely to be directly involved in DNA repair, replication or transcription in MCF7 cells. Even though *FAM111B*^*-/-*^ cells were proficient in their responses to DNA damage, we wondered whether their abnormal nuclear shape was associated with increased levels of endogenous DNA damage as is observed in mutants of the components of the nuclear lamina [49-51]. We screened the mutant cells for hallmarks of genomic instability. Quantification of micronuclei in the mutant cells (Figure 3D and E, Supplementary 2L) showed greater levels of these events in the absence of FAM111B protease compared to wild-type cells. Consistently, we could also observe higher frequencies of events upstream of micronuclei formation such as increased MIDAS foci (Figure 3F and G) and greater levels of BLM-coated DNA <u>u</u>ltra-fine <u>b</u>ridges (UFBs) in U2OS cells (Figure 3H and I). Re-introduction of wild-type but not protease dead (PD) or HFP mutation (Q430P, M1) variant into the *FAM111B*^*-/-*^ cells could rescue the normal levels of UFBs (Figure 3J and Supplementary Figure 3A). This confirmed that the observed phenotype is specific to the loss of FAM111B and that protease activity of FAM111B plays an essential role in maintaining low levels of BLM bridges in U2OS cells. We conclude that *FAM111B*-deficient cells even though proficient in DNA repair accumulate DNA damage and show hallmarks of genomic instability.

### Cells lacking FAM111B have short and abnormal telomeres

Beside the role that the nuclear pore complexes play in the responses to DNA damage they are also involved in the telomere lengthening and repair by recruiting telomeres to the nuclear envelope [45]. Interestingly, HFP patients develop pulmonary fibrosis which strongly correlates with short telomeres (reviewed in [1-5] and [33]). To test whether FAM111B may be required for telomere homeostasis, we looked at the length and condition of the telomeres in *FAM111B*^*-/-*^ cells.

Quantification of the telomeric FISH probe intensity in the metaphase spreads prepared from wild-type and mutant cells showed that cells lacking FAM111B protease indeed had greatly reduced telomeric DNA content (Figure 4A and B, similar phenotype in interphase cells was also observed, data not shown). Consistently, shorter telomeres in *FAM111B*^*-/-*^ U2OS cells recruited less of the shelterin component TRF2 compared to wild-type counterparts (Figure 4C and D). This phenotype was independent of passage number (Supplementary Figure 3B) and was not caused by off target effect of the CRISPR deletion method applied (Supplementary Figure 3C). Because pulmonary fibrosis observed in the FAM111B mutation carriers can be directly linked to short telomeres, we tested whether over-expression of HFP variants of FAM111B in U2OS cells had any effect on telomeres. Telomere shortening requires multiple rounds of division to be detectable and unfortunately, prolonged expression of FAM111B (wild-type or mutant but not protease dead [22], and our unpublished data) is toxic, preventing direct measurement of telomere length in extended experimental conditions. Therefore, we transiently over-expressed different variants of FAM111B in U2OS cells: wild-type, protease dead and HFP mutations, for no longer than 24h and instead quantified TRF2 recruitment to the telomeres. Surprisingly, wild-type and mutant protein but not protease dead variant of FAM111B induced loss of TRF2 intensity. This indicates that over-expression of FAM111B wild-type or mutant protein can interfere with the levels of telomere bound TRF2 (Figure 4E and Supplementary Figure 3F), and suggests that as well as a normally regulated proteolysis function, the cell cycle regulation of FAM111B is likely necessary for proper telomere function. Because only critically short telomeres are detrimental for cell viability and genome integrity [52], we looked whether *FAM111B*^*-/-*^ cells had any telomeric abnormalities beside short length. The same metaphase spread samples were used to determine levels of four phenotypes (Figure 5A-D): loss of telomere signal, telomere fusion, telomere heterogeneity and fragility. Both cell lines in the absence of FAM111B protease showed increased levels of telomere loss and fusion events (Figure 5A, B and E), whereas telomeric heterogeneity and fragility was specifically increased only in U2OS cells (MCF7 on average showed slight decrease, Figure 5C, D and E). Loss of FAM111B induced telomere shortening both in telomerase positive MCF7 cells and in U2OS cells which utilise alternative lengthening of telomeres (ALT), strongly indicating that FAM111B operates independently of these telomere lengthening mechanisms. Consistently with this, levels of C-circles in the *FAM111B*-deficient U2OS cells were comparable to wild-type (Supplementary Figure 3D) and MCF7 knockout cells showed wild-type like survival to telomere replication inhibitor TMPyP4 (Supplementary Figure 2K).

**Figure 4.**
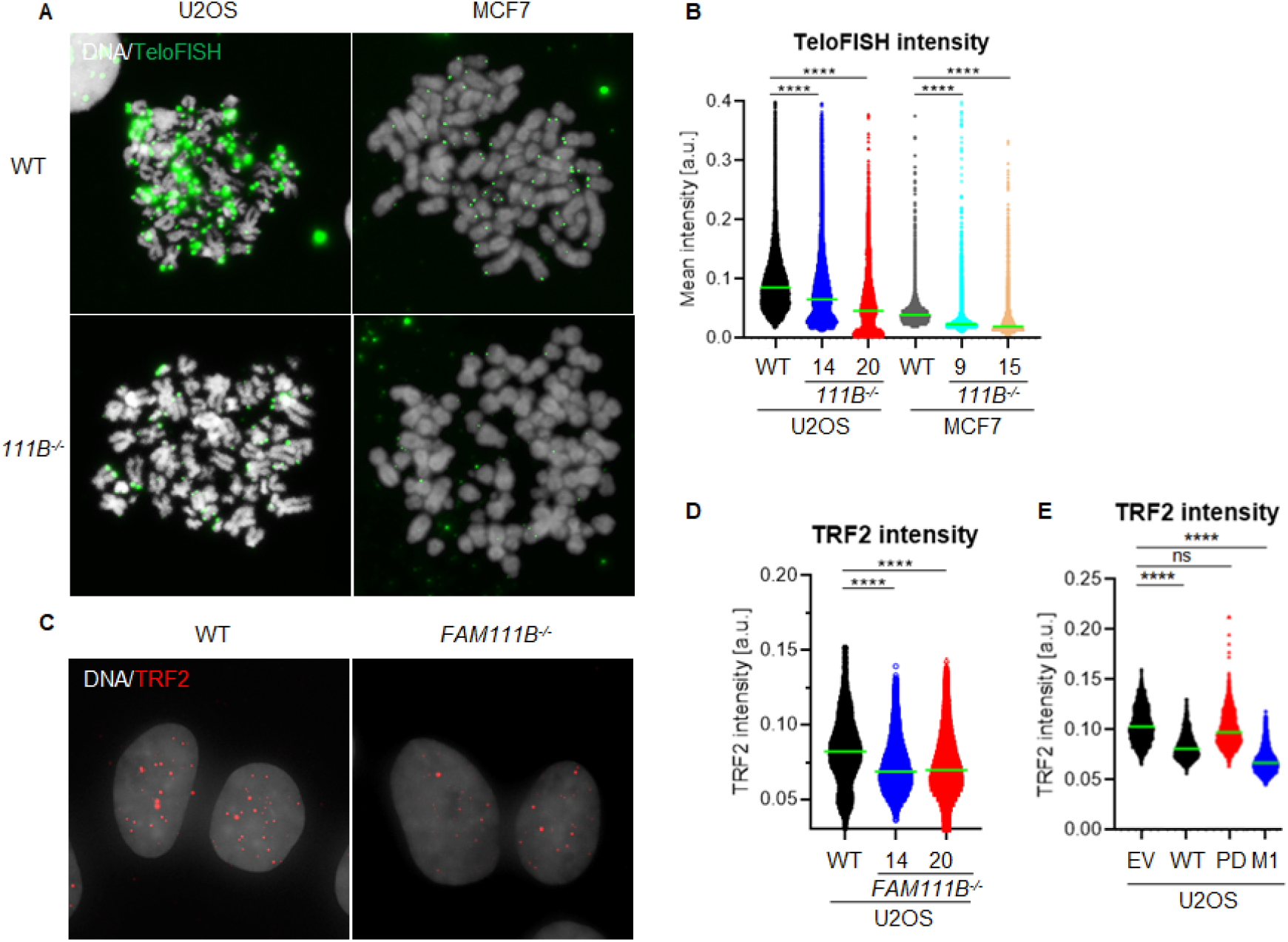
FAM111B-deficient cells have shorter telomeres. A) Images of chromosome spreads from wild-type (WT) and FAM111B knockout cells stained with fluorescent probe against telomeric repeats. B) Quantification of TeloFISH signal intensity. C) Images of wild-type (WT) and FAM111B^-/-^ U2OS cells stained with TRF2 antibodies. D) and E) Quantification of TRF2 foci intensity in U2OS wild-type (WT) and FAM111B negative cells or U2OS cells over-expressing FLAG-tagged FAM111B variants wild-type, protease-dead (PD) or HFP mutant (Q430P, M1), respectively.

**Figure 5.**
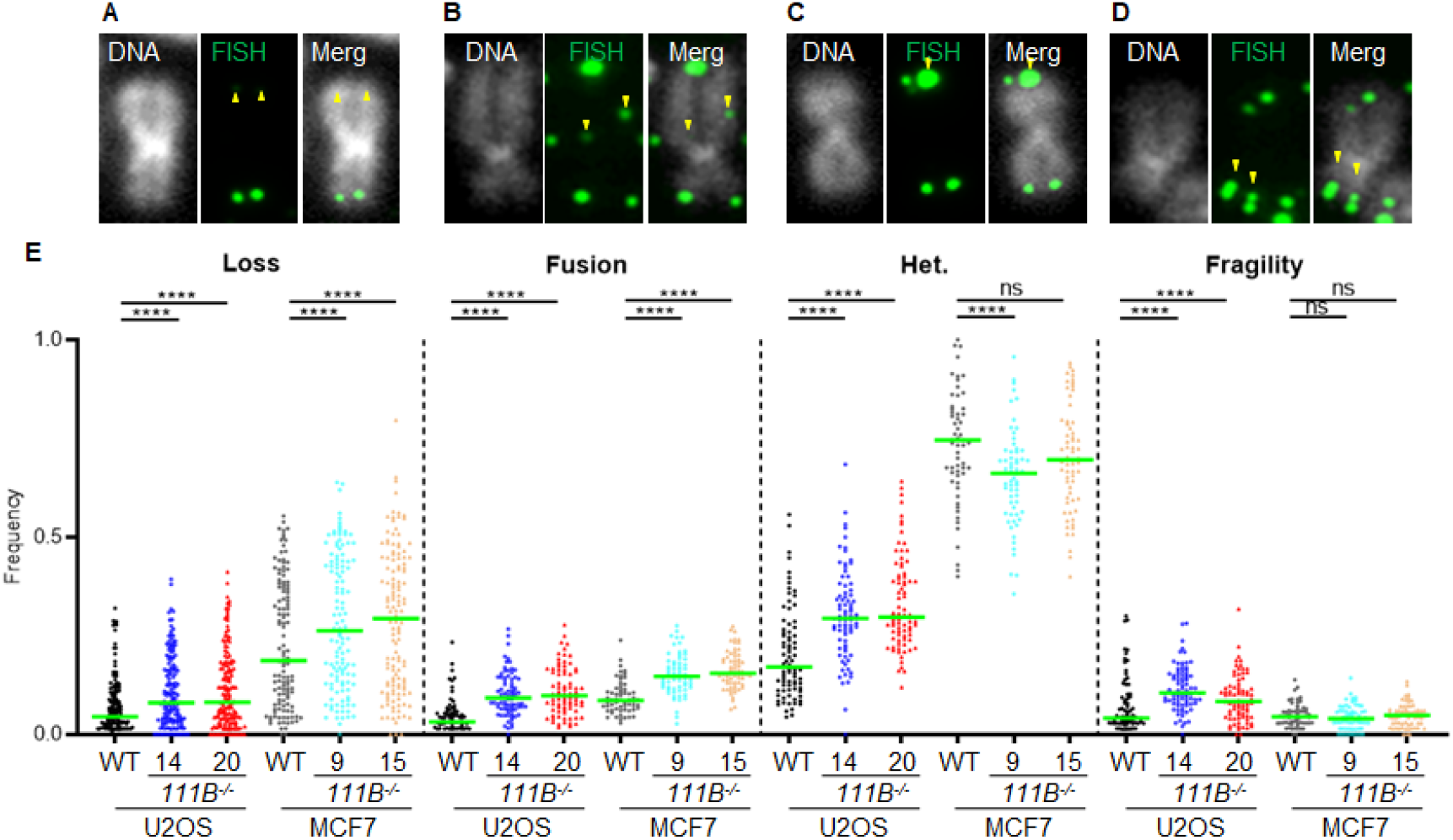
Loss of FAM111B induces telomere abnormalities. A) to D) Images of individual chromosomes stained with a fluorescent probe against telomeric repeats, showing telomeric abnormalities detected in FAM111B-deficient cells. These include telomere loss, fusion, heterogeneity and fragility. E) Quantification of telomeric phenotypes U2OS and MCF7 wild-type (WT) and FAM111B^-/-^.

## Discussion

Our analysis of FAM111B cellular functions showed that the protease is a non-essential, nuclear protein regulated by the cell cycle (Figure 1A and B). We discovered that a subpopulation of FAM111B localises to the nuclear periphery where it interacts with nuclear pore complexes (Figure 1C and D) but not with the components of the nuclear lamina such as lamins or emerin (Figure 1D). Mutant FAM111B is more frequently localised to the nuclear periphery and disease relevant variants accumulate in the insoluble fraction of the cell (Figure 2A-E). Loss of FAM111B induces nuclear shape abnormalities (Figure 3A-C) and genomic instability (micronuclei, MIDAS and UFBs) (Figure 3D-J). Interestingly, FAM111B is not required for normal responses to DNA damage or inhibition of DNA replication / transcription (Supplementary Figure 2A-J) but is needed to maintain normal telomere length in both telomerase- and ALT-positive cells. In the absence of FAM111B protease, telomeres are shorter and recruit less of the shelterin component TRF2 (Figure 4A-D). Many chromosomes in *FAM111B*-deficient cells showed no detectable telomeric DNA and frequently engaged in end-to-end fusion events (Figure 5A-E). This suggests that loss of FAM111B causes the formation of short telomeres. Similarly, cells over-expressing wild-type and HFP mutations (but not protease dead variant) have reduced TRF2 foci intensity (Figure 4E).

The FAM111B paralog protease FAM111A interacts with PCNA via a PIP-box motif and is required for removal of DNA-protein crosslinks [21]. In our hands *FAM111B*-deficient cells were no more sensitive to cis-platin, camptothecin, etoposide and formaldehyde that, besides damaging DNA, also induce protein-DNA crosslinks (Supplementary Figure D, E, F and J). This suggests that any role of FAM111B at replication forks may be different from FAM111A. 3D Genome organisation is dependent on robust interactions with the nuclear lamina and disruption of the nuclear envelope can inhibit DNA synthesis [53, 54]. Because FAM111 proteases interact with components of the nuclear pore complexes (this manuscript and [22, 55]), FAM111 proteases may influence DNA replication and transcription by targeting proteins at this specific nuclear localisation. Currently no substrate of FAM111B has been identified but at least one component of the nuclear pore complex (NUP62) has been recently described as a target of FAM111A [55]. Because of the many similarities between the two proteases, we hypothesise that FAM111B likely also target components of the NPC. In our current working model for FAM11B function this protease activity is needed in properly controlled fashion at NPCs to maintain normal telomeric structures. In the absence of FAM111B, or the presence of HFP-causing mutations, critically short telomeres induce DNA synthesis in mitosis (MIDAS), BLM bridges and micronuclei formation in the following G1 phase of the cell cycle (Figure 2D-J). Consistent with this model, lymphoblasts isolated from HFP patients showed spontaneous chromosome aberrations [56]. The cellular phenotypes described in this manuscript could explain why patients carrying FAM111B mutations suffer from pulmonary dysfunction as fibrosis of the lungs strongly correlates with short telomeres [34-38]. Interestingly, mutant FAM111B proteins including HFP associated mutations showed more prominent localisation to the nuclear periphery (Figure 2A-D) suggesting a potential mechanism of the disease pathology where deregulated FAM111B-dependent processing of yet unidentified target(s) at the proximity of nuclear pore complexes causes shorter telomeres. This in turn precipitates spontaneous DNA damage and affect cellular fitness resulting in threshold-specific deterioration of tissues such lungs, skin, muscles and tendons.

## Materials and methods

### Cell culture

Human osteosarcoma (U2OS), human breast cancer (MCF7) and human embryonic kidney cells (HEK293T) cells were cultured in Dulbeco’s Modified Eagle Medium (DMEM, Sigma) supplemented with 10% fetal bovine serum (Sigma), 1% glutamate (Sigma) and 1% penicillin/streptomycin (Sigma) at 37°C in humidified 5% CO_2_ atmosphere. For conditional induction of protein expression 1μg/ml of doxycycline (Sigma) was added to the culture media and incubated for 16-24h.

### Gene and CRISPR construct cloning

All gene cloning was performed using Gateway® system (Invitrogen) unless stated otherwise.

Briefly, human *FAM111B* cDNA (FW: GGGGACAAGTTTGTACAAAAAAGCAGG CTTCAATTCCATGAAGACTGAAGAAAAC and RV: GGGGACCACTTTGTACAAGAAAGCT GGGTCCTAACATTCCATGGGTTCAATC) was amplified using high fidelity Q5 DNA polymerase (New England Biolabs), cDNA template (kind gift from Dr Nicola Burgess-Brown) and cloned into pDONR-ZEO (Invitrogen) vector to obtain pDONR-ZEO-FAM111B-WT (wild-type). Protease dead (H490A, FW: CCACCATAAGATGTACAACAGCTCGACAGGTGAAAA TATAACCAC and RV: GTGGTTATATTTTCACCTGTCGAGCTGTTGTACATCTTATGG TGG), patient mutations (Q430P, FW: GAAGTCCTTTTTATATATGTTGAATGGCTTAGCT GGTGAAAGATTCATTC and RV: GAATGAATCTTTCACCAGCTAAGCCATTCAACATATAT AAAAAGGACTTC; T625N, FW: TCTGATAGGAAACTTCTTTGGTTAAACATACAGTATACA TTACTGGT and RV: ACCAGTAATGTATACTGTATGTTTAACCAAAGAAGTTTCCTATC AGA) and ΔED (Δ279-386, FW: ACGTTTGATAATTCTTTAAGAGATCTTTCTGTTGTAATGC TTTTTTTTTTGAAATGTCC and RV: GGACATTTCAAAAAAAAAAGCATTACAACAGAAAG ATCTCTTAAAGAATTATCAAACGT) variant were constructed by site directed mutagenesis using pDONR-ZEO-FAM111B-WT as a template and Q5 polymerase. Double mutant (H490A/T625N) was generated by sub-cloning T625N mutation from pDONR-ZEO-FAM111B-T625N into pDONR-ZEO-FAM111B-H490A by restriction digest and sticky end ligation. *FAM111B* cDNAs were then transferred from pDONR-ZEO to pHAGE-(N)-HA-FLAG-(PURO) vector for constitutive expression or to pSIN3A-(C)-HA-(PURO) vector for conditional expression (TET^ON^) (kind gifts from Prof Ross Chapmann). *FAM111B* targeting constructs (gRNA #1: CACCGGAAGGAGCCTTATGCAAGGA and AAACTCCTTGCATAA GGCTCCTTCC and gRNA #2 (CACCGTCCAGATACTTCATCCA CCA and AAACTGGTG GATGAAGTATCTGGAC) were cloned into pX330-(PURO) (kind gift from Prof Ross Chapmann) using previously published protocol [57].

### Antibodies

Primary antibodies were as follows: mouse anti-human BLM (sc-365753, Santa Cruz Biotechnology) at 1:250 (IF), mouse anti-BrdU (347580, Becton Dickinson) at 1:20 (Flow cytometry), rabbit anti-human FAM111B (HPA038637, Atlas Antibodies) at 1:1000 (IB and IF), mouse anti-human Histone H3 (10799, Abcam) at 1:1000 (IB), mouse anti-human Lamin A/C (sc-376248, Santa Cruz Biotechnology) at 1:1000 (IB and IF), rabbit anti-human Lamin B1 (12987-1-AP, Proteintech Group) at 1:1000 (IB and IF), rabbit anti-human NUPL2 (16587-1-AP, Proteintech Group) at 1:1000 (IB), mouse anti-human PCNA (sc-56, Santa Cruz Biotechnology) at 1:1000 (IB and IF), rabbit anti-human SEC13 (15397-1-AP, Proteintech Group) at 1:1000 (IB), mouse anti-human TRF2 (NB100-56506, Novus Biologicals) at 1:1000 (IF), mouse anti-human Tubulin (T6074, Sigma) at 1:10 000 (WB) or 1:1000 (IF). Secondary antibodies were as follows: goat anti-rabbit Alexa Fluor 488 (A32731, ThermoFisher Scientific) at 1:1000 (IF), donkey anti-mouse Alexa Fluor 555 (A32773, ThermoFisher Scientific) at 1:1000 (IF), goat anti-rabbit HRP (31430, ThermoFisher Scientific) at 1:25 000 (IB) and donkey anti-mouse HRP (31458, ThermoFisher Scientific) at 1:25 000 (IB).

### Microscopy and immuno-fluorescence

U2OS or MCF7 cells were seeded at 2×10^5^ onto glass cover slips in 6 well plates. 48h post-seeding cells were fixed with 4% formaldehyde in PBS for 10min at room temperature, washed three times with PBS (3min each), permeabilised with CSK buffer for 5min at room temperature (10mM PIPES-KOH pH6.8, 100mM NaCl, 300mM sucrose, 1mM EGTA, 1mM MgCl_2_ and 1mM DTT) supplemented with protease and phosphatise inhibitors and again washed three times with PBS. For detergent extraction experiments, cells were first exposed to CSK buffer for 5min at room temperature washed three times with PBS (3min each) and then fixed with 4% formaldehyde for 10min at room temperature, followed by washes with PBS (3min each). Cover slips were then blocked with 3% BSA/PBS for 30min at room temperature, incubated with primary antibodies in the blocking buffer for 1h at room temperature and washed three times with PBS. This was followed by the incubation of cover slips in secondary antibodies diluted in the blocking buffer for 1h at room temperature, three washes with PBS and mounting the cover slips onto glass slides with Vectashield+DAPI (H-1200-10, 2BeScientific). For staining of ultra-fine DNA bridges (UFBs), 2×10^5^ cells was seeded onto cover slips and 24h later cells were exposed to 2mM thymidine for 16h. The next day, the nucleotide was washed away (three times with warm media) to release cells from the block and trapped in pro-metaphase for 3h at 9h post-thymidine removal with the addition of nocodazole at 200ng/ml. Microtubule poison was then washed away (three times with warm fresh media) and cells harvested 30-45min after release from the nocodazole block. For the staining of UFBs in the *FAM111B* rescue cell lines, doxcycline was added exactly 24h prior to 30min release mark from the nocodazole block. For detection of EdU foci in prometaphase cells (MIDAS), U2OS cells were seeded the same way as in for analysis of UFBs, however; cells were not released from the nocodazole block, labelled for last 30min with 10μM EdU and fixed with 4% formaldehyde for 10min at room temperature. EdU detection was performed according to the manufacturer’s instructions (Invitrogen). Analysis was performed using wide-field Leica DM4000 or AyraScan 2 (Zeiss) microscopes.

### Cell transfections

For transient transfections of U2OS or HEK293T cells, 70-80% confluent cultures in 6 well plates or 10 cm dishes were transfected using 3µg or 18µg of linear PEI (Sigma), 0.2ml or 1ml OPTI-MEM medium (Life Technologies) and 1µg or 6µg of DNA, respectively. Cells were analyzed by microscopy or harvested 24h post-transfection. To obtain stably expressing U2OS clones, 70-80% confluent cultures in 10cm dishes were transfected as above and 24h post-transfection cells were exposed to Puromycin (Invitrogen) at 0.5µg/ml. After approximately 2-3 weeks, single colonies were collected by trypsinisation and expanded.

### Generation of FAM111B-deficient clones

As described in the section “Cell transfections”, pX330-(PURO)-FAM111B gRNA #1 and #2 constructs were delivered to U2OS and MCF7 by transfections with linear PEI. After 24h post-transfection, cells were pulsed with puromycin at 1ug/ml for 72h. Approximately 2-3 weeks later, single colonies were collected by trypsinisation and expanded in 24 and then 6 well plates. When clones reached 90% confluency in 6 well plates, half of the cultures was frozen and the remaining cells harvested and analysed by immune-blotting with antibodies against FAM111B and Tubulin as a loading control. Clones negative for FAM111B expression were revived from liquid nitrogen and expanded.

### Cell fractionation

Approximately 2×10^6^ of U2OS cells (either transiently transfected or not) was used for these experiments. Cells were washed once with ice cold PBS buffer, split in half (fractionation and whole lysate sets) and spun at 300g for 3min. Fractionation set was re-suspended in 100μl of buffer A (10mM HEPES pH7.4, 10mM KCl, 1.5mM MgCl_2_, 340mM sucrose, 10% glycerol and 1mM DTT) supplemented with protease and phosphatase inhibitors (Sigma). 2μl of Triton X-100 (from 5% stock) was then added to the fractionation set, mixed for 3s using vortex (highest speed) and placed on ice for 5min. Lysates were then spun at 1300g for 4min at 4°C, 80μl of the supernatant was collected as soluble fraction and mixed with 20μl of 5x Laemmli buffer. Remaining pellet was gently washed with 1ml of buffer A (pellet was not disturbed), spun again and re-suspended in buffer C (50mM Tris-HCl pH7.4, 9M Urea and 150mM 2-mercaptoethanol), thoroughly sonicated and spun at 21 000g.

Similarly, 80μl of the supernatant was collected as insoluble fraction and mixed with 20μl of 5x Laemmli buffer. The whole lysate set was re-suspended in buffer C, sonicated and spun at 21 000g for 15min at 4°C. Again, 80μl of the supernatant was collected as whole cell lysate sample and mixed with 20μl of 5x Laemmli buffer. All samples were then boiled for 5min at 95°C and analysed by immuno-blotting with the indicated antibodies.

### C-circle assay

DNA was extracted from cells (10^6^ cells per sample) using the QIAGEN Core B kit and re-suspended in 20mM Tris-HCl. 30 ng of genomic DNA was amplified in a PCR reaction containing φ29 polymerase, 1% Tween-20, 200 µg/mL BSA and dTTP, dGTP and dATP for 8 hours at 30°C followed by 20 minutes at 65°C. PCR reactions were run with and without φ29 polymerase to ensure the signal was specific for rolling-circle amplification products. Amplified samples were then blotted onto Zeta-Probe membrane (Bio-Rad) using a slot blotter. DNA was crosslinked to the membrane with a UVA Stratalinker 2400 and membranes were then soaked in PerfectHyb Plus (Sigma Aldrich) for 20 minutes at room temperature. A 3’ Digitonin (DIG) tagged [CCCATT]_5_ oligonucleotide was then diluted in PerfectHyb to a final concentration of 40 nM in 20mL hybridisation buffer, and was hybridised with the membrane for 2 hours at 37°C. Following hybridisation, membranes were briefly washed twice with wash buffer (0.1M Maleic acid, 3M NaCl; 0.1% Tween 20 adjusted to pH 7.5), before blocking for 30 minutes and probing with a DIG antibody for 30 minutes. Finally, membranes were washed 3 times with wash buffer before placing into a cassette with CDP-Star solution. Blots were then developed onto Amersham Hyperfilm ECL film.

### Growth curves and cell survival assay

For growth curves, MCF7 and U2OS cells were seeded in duplicate in 96 well plates at 2500 cells per well (including no cells blank). Each day, separate duplicate wells were incubated with DMEM Fluorobrite Media (A1896701, GIBCO) supplemented with 10% fetal bovine serum, 1% glutamate, 1% penicillin/streptomycin and 10μg/ml resazurin for 2h. After the incubation time, fluorescence in the appropriate wells was read using microplate reader (CLARIOStar^Plus^, BMG LABTECH) with the following settings: excitation λ = 530nm, emission λ = 590nm. For the cell survival assays, 24h post-seeding old media was replaced with fresh media containing the indicated drugs at different doses and cells incubated for another 72h. On the final day the survival plates were assayed identically as growth curve plates. Cell growth was expressed as a relative % of wild-type cells whereas survival of the cells was expressed as relative % of the corresponding untreated samples.

### Pull-down assays

For pull-down of FLAG-tagged proteins, transiently transfected HEK293T (single 10cm dish) were lysed in pull-down buffer (20mM HEPES pH 7.4, 150mM KCl, 10% glycerol, 2mM MgCl2, 0.5mM DTT, 0.5% NP-40) supplemented with protease and phosphatase inhibitors (Sigma) as well as benzonase (250 U; Sigma) for 1 h at 4°C on a rotating wheel. Lysates were then spun at 21,000 g for 15 min at 4°C and protein concentration determined using Bradford assay (Sigma). Approximately 3-4mg of total cell extract was mixed with 20μl of M2-anti-FLAG agarose beads (bed volume) previously washed once with 1ml of pull-down buffer and incubated for 2h at 4°C on a rotating wheel. After the incubation, beads were spun (30s at 2700g at 4°C), washed 5 times with the pull-down buffer (at the second wash beads were transferred to a fresh tube) and eluted with 30μl of 3xFLAG peptide at 400μg/ml for 30min at 4°C shaking. Elutions were then mixed with 5x Laemmli buffer, boiled for 5min at 95°C and analysed by immuno-blotting with the indicated antibodies.

### Flow cytometry

Approximately, 10^6^ of U2OS or MCF7 cells were pulsed with 50μM BrdU for 5min, harvested by trypsinisation and re-suspended in 1ml of PBS. 2ml of -20°C absolute ethanol was then added dropwise to the cells while gently vortexing. Cells were fixed for at least 30 min at room temperature, spun for 5min at 500g and ethanol removed by two washes with PBS. To reveal BrdU epitopes, samples were treated with 1ml of 2M HCl/ 0.5 % Triton X-100 solution for 30min at room temperature, followed by acid neutralisation with 1ml of 0.1M sodium tetraborate pH8.5. Cells were then blocked with 1% BSA/0.5% Triton X-100 in PBS for 30min at room temperature, counted and brought to concentration of 10^6^ cells / ml. 20μl of the anti-Brdu antibody was added to the cells and incubated for 1h at room temperature, followed by three washes with 1% BSA/0.5% Triton X-100 in PBS and incubation in 100μl of anti-mouse Alexa 488 in 1% BSA/0.5% Triton X-100 for 1h at room temperature. Samples where again washed three times with 1% BSA/0.5% Triton X-100 in PBS and treated with RNase A at 100 µg/ml and propidium iodide at 40 µg/ml in PBS for 30 min at room temperature. Analysis of the samples was performed using SH800z Cell Sorter instrument (SONY).

### Fluorescent in-situ hybridisation

Chromosome spreads were prepared with standard techniques. Briefly, cells were incubated with KaryomaxColcemid (Thermo Fisher Scientific) 50 ng/mL for 3 h. Following a swelling step in hypotonic (Buffered Hypotonic Solution, Genial Helix), the cells were fixed twice in Carnoy’s fixative (methanol:acetic acid 3:1). The cells suspension was dropped onto clean slides, and air-dried.

The copy number and position of the *FAM111B* locus in the different cell lines was analysed by FISH with fosmid G248P81352D4 (WI2-0409G07, hg19 chr11:58,859,240-58,898,785) mapping to *FAM111B* genomic position, and two BAC constructs flanking the region, RP11-100N3 (hg19,chr11:56481672-56641043) and RP11-286N22, hg19 chr11:61,197,622-61,249,870).The probes were labelled by nick translation, using a commercial kit (Abbott Molecular Nick Translation Kit) incorporating ChromaTide Alexa Fluor 488-5-dUTP (Thermo Fisher Scientific), ChromaTide Alexa Fluor 594-5-dUTP (Thermo Fisher Scientific), or biotin-16-dUTP (Sigma). The probes were re-suspended in hybridization buffer (50% formamide, 10% dextran sulfate, 2× SSC) at 10 ng/μL, in the presence of a 10× excess of unlabeled human Cot1 DNA (Sigma Aldrich). The cellular DNA was denatured in NaOH 0.07M. The probe DNA was denatured by incubating it at 85°C and allowed to renature at 37°C for 30 minutes. The hybridisation was carried out overnight at 37°C. Following post-hybridisation washes in 0.1xSSC at 60°C, the biotinylated probe was visualised using Avidin Cy5 (ThermoFisher Scientific). A minimum of 25 images per cell line were acquired and analysed with the Leica Cytovision software, on an Olympus) BX-51 epifluorescence microscope equipped with a JAI CVM4+ progressive-scan 24 fps black and white fluorescence CCD camera.

The TeloFISH was carried out using the Telomere PNA FISH Kit/Cy3 (Dako), following the manufacturer instructions. A minimum of 30 metaphases per cell line were acquired, using a Leica DM6B microscope for epifluorescence, equipped with a DFC 9000Gt B&W fluorescence CCD camera, and operated via the Leica LASX software.

### Mass spectrometry

Following the FLAG-IP, elutions were denatured in 200μl 8M urea (4.8g / 10ml) in 100mM TEAB for 30 min at RT, reduced with 10mM TCEP, for 30 min at RT, alkylated with 50mM iodoacetamide for 30 min at RT in the dark and finally diluted down to 1.5M urea with 50mM TEAB. Protein digestion was performed using 1.5 ng of trypsin in 50mM TEAB over night at 37C, followed by clean up with C18 column (PepMapC18; 300µm x 5mm, 5µm particle size, Thermo Fischer) using solvent A (0.1% Formic Acid in water) at a pressure of 60 bar and separated on an Ultimate 3000 UHPLC system (Thermo Fischer Scientific) coupled to a QExactive mass spectrometer (Thermo Fischer Scientific). The peptides were separated on an Easy Spray PepMap RSLC column (75µm i.d. x 2µm x 50mm, 100 Å, Thermo Fisher) and then electrosprayed directly into an QExactive mass spectrometer (Thermo Fischer Scientific) through an EASY-Spray nano-electrospray ion source (Thermo Fischer Scientific) using a linear gradient (length: 60 minutes, 5% to 35% solvent B (0.1% formic acid in acetonitrile and 5% dimethyl sulfoxide), flow rate: 250 nL/min). The raw data was acquired on the mass spectrometer in a data-dependent mode (DDA). Full scan MS spectra were acquired in the Orbitrap (scan range 380-1800 m/z, resolution 70000, AGC target 3e6, maximum injection time 100 ms). After the MS scans, the 15 most intense peaks were selected for HCD fragmentation at 28% of normalised collision energy. HCD spectra were also acquired in the Orbitrap (resolution 17500, AGC target 1e5, maximum injection time 128 ms) with first fixed mass at 100 m/z. Progenesis QI (Waters) based label-free quantitation results were imported into Perseus 1.5.2.463.

Quantitative data was log2 transformed and normalized by median subtraction and missing values were imputed based on normal distribution.

### Image analysis and statistics

Image analysis was performed using CellProfiler v. 4.2.1 software and home design pipelines. Briefly, for the measurements of TeloFISH foci intensities, individual chromosome objects were identified using Watershed algorithm and DAPI channel. Neighbouring nuclei were removed using twostep process: step one - Measure Object Size and step two Filter Objects by size (nuclei > chromosomes). Individual chromosomes were then used to mask TeloFISH channel (Mask Image) and telomers identified using Identify Primary Objects algorithm. Finally, intensity of individual TeloFISH signals was determined with Measure Object Intensity algorithm. Similarly, for the TRF2 foci intensity measurements, nuclei were identified in DAPI channel (Identify Primary Objects) and used as a mask (Mask Image) to identify nuclear TRF2 objects coming from telomeres (Identify Primary Objects). Intensity of TRF2 objects was then measured using Measure Object Intensity algorithm.

For the statistical calculations, Graphpad Prism 9 software was used. Briefly, if the population of interest followed the normal distribution parametric tests such as One-Way Anova were used, whereas if the populations did not follow the Gausian distribution non-parimetric test such as Kruskal-Wallis were used to determine whether the observed differences between sample groups were significant.

---

**Supplementary Figure 1.**
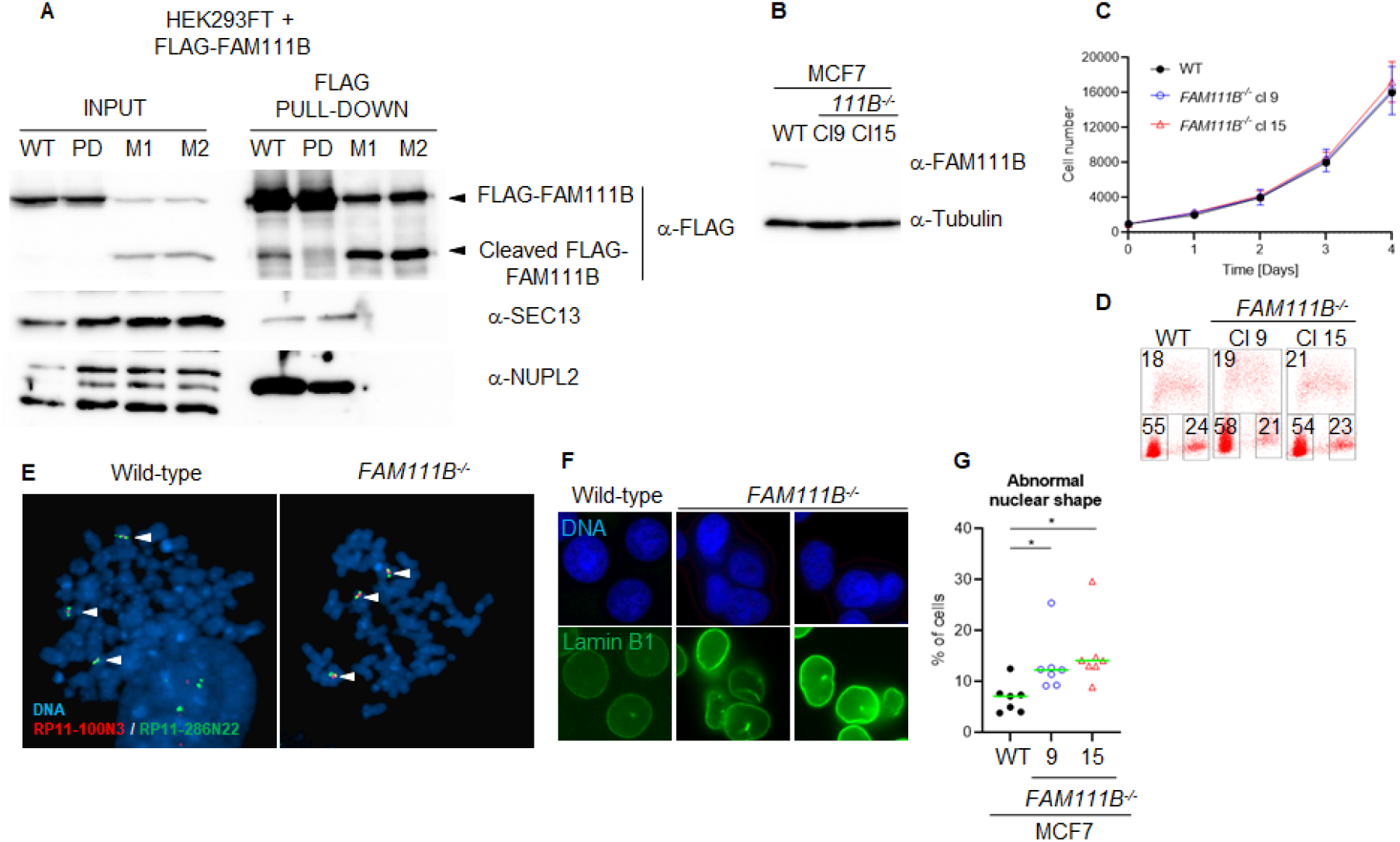
FAM111B interacts with the components of the nuclear pore complex. A) Immuno-blots of FLAG-FAM111B pull-downs showing complex formation between FAM111B wild-type (WT), protease dead (PD), HFP mutant (Q430P, M1) and subunits of the nuclear pore complex SEC13 and NUPL2. B) Immuno-blots showing loss of FAM111B in the MCF7 knockout cell lines. C) Growth curve of wild-type (WT) and FAM111B-deficient MCF7 clones. D) Flow cytometry analysis of the cell cycle distribution in MCF7 wild-type and two FAM111B knockout clones. E) Images of chromosome spreads stained with fluorescent probes specific to FAM111B and control loci. F) Images of MCF7 wild-type and FAM111B-deficient cells stained with Lamin B1. G) Quantification of abnormally shaped nuclei in wild-type and FAM111B knockout cells.

**Supplementary Figure 2.**
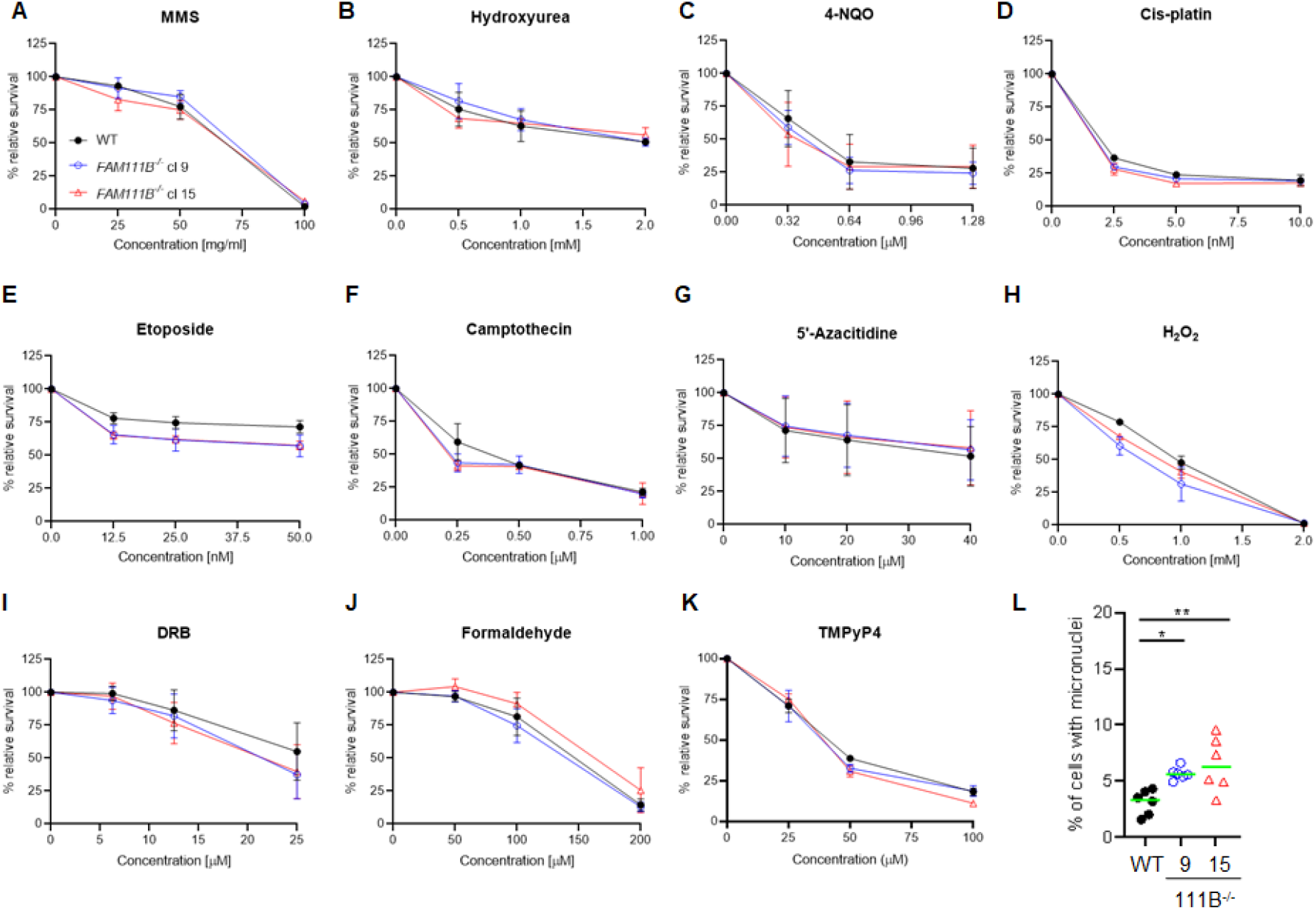
FAM111B-deficient cells show wild-type like response to DNA damage and DNA replication / transcription perturbations. A) to K) survival of MCF7 wild-type (WT) and FAM111B knockout cells after chronic exposure to the indicated drugs. Error bars show standard error, N=3. L) Quantification of micronuclei in MCF7 wild-type and FAM111B negative cells.

**Supplementary Figure 3.**
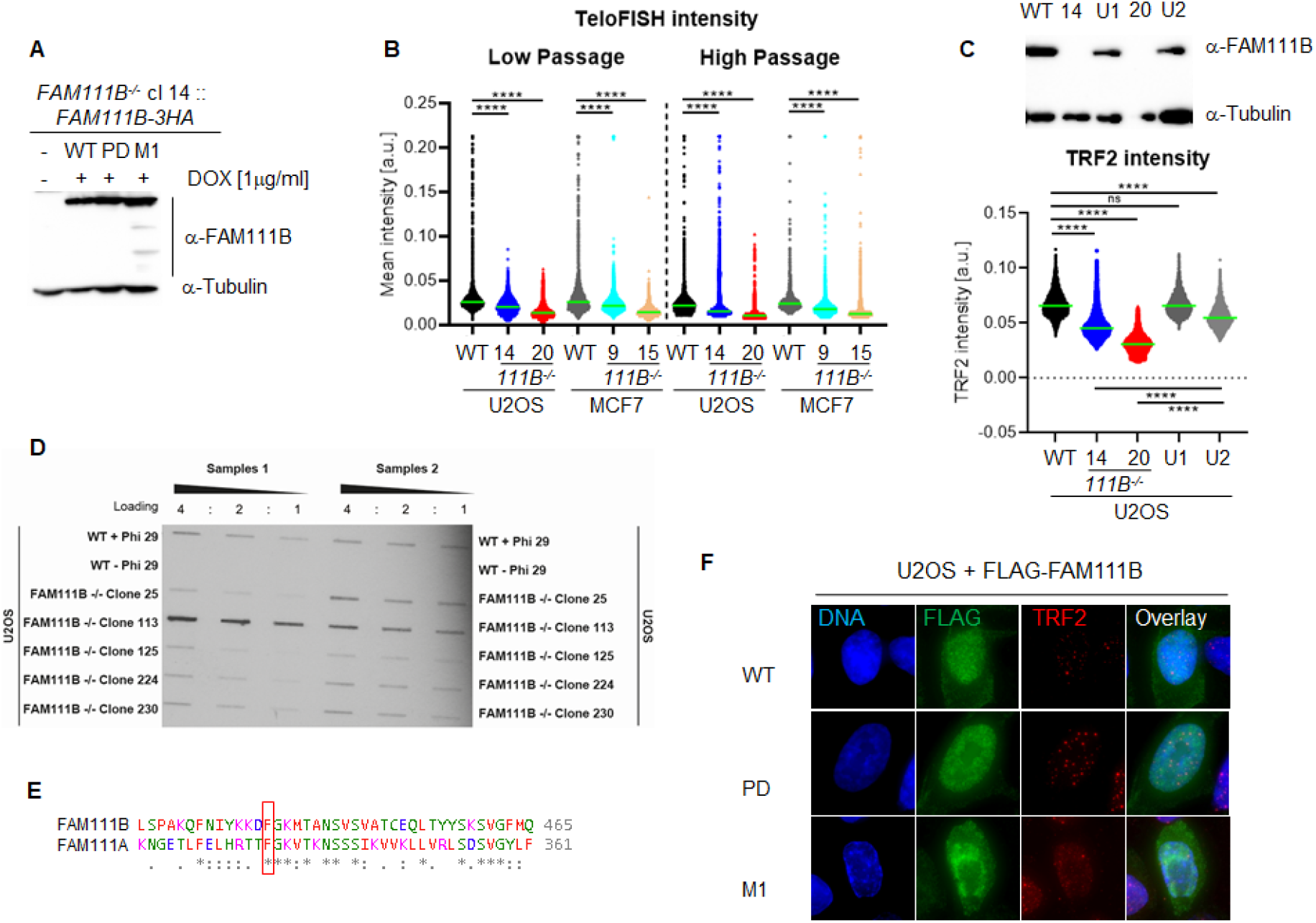
Loss of FAM111B induces short telomeres and interferes with TRF2 recruitment. A) Immuno-blot of U2OS FAM111B^-/-^ cells in which FAM111B wild-type (WT), protease dead (PD) or HFP variant (Q430P, M1) where re-introduced under the control of doxycycline (DOX) inducible promoter. B) Quantification of TeloFISH signal intensity in U2OS and MCF7 cells at low and high passage numbers. C) Immuno-blot of FAM111B levels in targeted and untargeted U2OS (top panel). Quantification of TRF2 foci intensity in these clones (bottom panel). D) Southern slot blot showing levels of C-circles in multiple U2OS FAM111B-deficient clones. E) Protein sequence alignment of FAM111B and FAM111A surrounding the described auto-cleavage site in FAM111A protease [21]. F) Images showing U2OS cells transiently expressing FLAG-FAM111B wild-type, protease dead and HFP variants stained with FLAG and TRF2 antibodies.

***Movie 1A. Localisation of FAM111B to the nuclear periphery***. *Three-dimensional reconstitution of FAM111B staining in U2oS cells (DAPI staining DNA in Blue, Antibodies staining of FAM111B in Green and Lamin A/C in Orange) U2OS cells. Movie shows different staining and masks:*

*Turn 1 – DNA* | *total FAM111B* | *Lamin A/C*

*Turn 2 - DNA*

*Turn 3 - Lamin A/C*

*Turn 4 - Total FAM111B*

*Turn 5 – DNA* | *Nuclear FAM111B*

*Turn 6 – DNA* | *Lamin A/C masked FAM111B*

*Turn 7 – Lamin A/C* | *Lamin A/C masked FAM111B*.

***Movie 1B. Localisation of FAM111B to the nuclear periphery***. *Three dimensional reconstitution of FAM111B staining in U2OS cells (DAPI staining in Blue and antibodies staining of FAM111B in Green) showing only FAM111B signal masked with Lamin A/C (not shown)*.

## Notes

### Competing Interest Statement

The authors have declared no competing interest.

